# Evaluating the effects of alcohol and tobacco use on cardiovascular disease using multivariable Mendelian randomization

**DOI:** 10.1101/757146

**Authors:** Daniel B. Rosoff, George Davey Smith, Nehal Mehta, Toni-Kim Clarke, Falk W. Lohoff

## Abstract

Alcohol and tobacco use, two major modifiable risk factors for cardiovascular disease (CVD), are often consumed together. Using large publicly available genome-wide association studies (results from > 940,000 participants), we conducted two-sample multivariable Mendelian randomization (MR) to simultaneously assess the independent effects of alcohol and tobacco use on CVD risk factors and events. We found genetic instruments associated with increased alcohol use, controlling for tobacco use, associated with increased high-density-lipoprotein-cholesterol (HDL-C), decreased triglycerides, but not with coronary heart disease (CHD), myocardial infarction (MI), nor stroke; and instruments for increased tobacco use, controlling for alcohol use, associated with decreased HDL-C, increased triglycerides, and increased risk of CHD and MI. Exploratory analysis found associations with HDL-C, LDL-C, and intermediate-density-lipoprotein metabolites. Consistency of results across complementary methods accommodating different MR assumptions strengthened causal inference, providing strong genetic evidence for the causal effects of modifiable lifestyle risk factors on CVD risk.

## INTRODUCTION

Cardiovascular disease (CVD) is a leading cause of mortality and morbidity with an estimated annual 17.9 million deaths worldwide^1^. Evidence shows that addressing modifiable lifestyle factors can prevent most CVDs^2^. Two highly prevalent and frequently co-occurring risk factors, alcohol consumption and smoking (affecting 2 billion and 1.1 billion people worldwide, respectively^2,3^) are associated with a considerable proportion of CVD mortality. For example, one in ten CVD deaths are attributable to smoking^4^, and observational studies strongly suggest that smoking increases the incidence of myocardial infarction (MI) and coronary heart disease (CHD)^5^.

Using novel epidemiological methods, the Global Burden of Diseases, Injuries, and Risk Factors Study (GBD) recently reported that any level of alcohol monotonically increases risk for many health-related harms, including cancer (GBD)^6^; however, considerable debate remains regarding the association between alcohol consumption and CVD risk ^7,8^

Observational studies show a complex relationship between alcohol consumption and CVDs with some studies reporting light-to-moderate alcohol use associated with moderately reduced risk of MI^9^ and coronary heart disease^10,11^, while heavier alcohol consumption and binge drinking are associated with increased stroke, MI, and CHD ^11,12^. Meta-analyses and short-term trials suggest alcohol consumption is associated with CVD risk factors including increased high-density lipoprotein-cholesterol (HDL-C) ^13^; however, the association with low-density lipoprotein cholesterol (LDL-C) and triglycerides (TRG) are less clear ^14^. Variation in size, density, concentrations and composition of circulating lipids, which traditional measures are unable to distinguish, are thought to have contrasting effects to CVD risk^15^. Metabolomics provides a detailed view of systemic metabolism, and the physiological effect of alcohol consumption extends across multiple metabolic pathways^7^ and has been associated with molecular dysregulation conferring both reduced and increased CVD risk^7^.

Despite its public health importance ^13^, nearly all human data evaluating the alcohol-CVD relationship is from conventional epidemiological studies^14^, and thus, the causal effect of alcohol consumption patterns on CVD risk factors and outcomes has not been fully elucidated. Triangulating evidence from different study designs is needed to improve our understanding of the alcohol-CVD relationship ^16^. However, observational studies are subject to potential confounding and reverse causation, which makes causal inferences difficult^17^ and many of the interventional studies or clinical trials examining the association between alcohol and CVD risk factors were small, failed to randomize different amounts of alcohol, and only observed short-term effects^13,18^. Further, the recent Moderate Alcohol and Cardiovascular Health (MACH) study, which was a large multicenter, worldwide, randomized clinical trial of ∼15 gm of alcohol daily versus abstention, has been stopped, due to concerns about study design and future credibility^19^.

One alternative strategy to investigate potential causal inference between an exposure and an outcome, in particular when randomized control trials (RCTs) are not feasible, practical, or ethical, would be Mendelian randomization analysis (MR)^20^. MR analyses can be conceptualized as natural RCTs^21^. MR utilizes randomly inherited genetic variants, established before outcome onset and therefore relatively independent of confounders, as instruments for an exposure to assess the causal effect of the risk factor exposure on a health outcome of interest^22^. Where it may be difficult to find a genetic variant uniquely associated with a risk factor, MR has been extended to multivariable MR (MVMR) using genetic variants associated with multiple risk factors—alcohol and tobacco use in this study—to simultaneously and consistently estimate the effect of each of the risk factors on the outcome(s) of interest^23,24^. Assumptions underlying single and multivariable MR analysis require sufficient statistical power to perform the required sensitivity analyses^22^. The two-sample single- and multivariable MR analysis used in this study—using summary-level data from separate genome-wide association studies (GWASs) for exposures and outcomes—allows for the use of complementary MR methods capable of investigating these assumptions ^25^.

Previous one-sample MR studies using genetic variation in either the acetylaldehyde dehydrogenase (*ALDH*) or alcohol dehydrogenase (*ADH*) genes found alcohol associated with increased HDL-C levels^8,18,26^. MR studies have shown that alcohol consumption reduces triglycerides and CHD risk^8^. A recent MR study using 512,715 Chinese adults from the Kadoori Biobank found alcohol increased stroke risk^27^, but had no effect on MI, possibly due to low MI prevalence in the sample^27^. Given the complexity of the relationship between alcohol and CVDs, average alcohol consumption may not be sufficient to evaluate the risk conferred for CVDs^28^. There is observational evidence for a link with approximately 85% of smokers drinking alcohol^29^, and drinkers are 75% more likely than abstainers to smoke^30^, as well as a strong genetic correlation^31,32^. Thus, failure to account for smoking in previous single variable MR analyses (SVMR) of alcohol and CVDs may have produced biased and inconsistent estimates.

Using a two-sample MR design (Fig. 1), we created instruments for alcohol and tobacco use using summary statistics from publicly available, large-scale genome-wide association studies (GWAS)^32^ (Table 1 and Supplementary Table 1) to evaluate the potential causal effects thereof on cardiovascular disease risk factors and disease events. Given the strong association between drinking and smoking^33^, we conducted multivariable MR analyses to estimate the direct effect of each of exposures—alcohol and tobacco use—controlled for the other—on our cardiovascular outcomes of interest. We compared these direct independent effects to the total effects estimated by single variable MR analysis. We also conducted exploratory multivariable MR analyses examining how alcohol and tobacco use effected changes in the metabolomic profile: assessing the detailed metabolomic effects of alcohol and tobacco use could inform our understanding of association thereof with lipid CVD biomarkers. Complementary MR techniques—inverse-variance weighted MR and MR-Egger—were used as sensitivity analyses to test the robustness of all results.

**Table 1.** Phenotype sources.

| Phenotype | Source | First Author (Year) | Sample Size | N Cases | N Controls | Population | Outcome Units |
| --- | --- | --- | --- | --- | --- | --- | --- |
| **EXPOSURES:** |  |  |  |  |  |  |  |
| Alcohol consumption - Drinks Per Week (DPW) | SSGAC (UKB) | Karlsson Linnér (2019) | 414343 | NA | NA | European | Continuous |
| Ever Smoker Status (ES) | SSGAC (UKB) | Karlsson Linnér (2019) | 518633 | NA | NA | European | Binary |
| **PRIMARY OUTCOMES:** |  |  |  |  |  |  |  |
| Cardiovascular Disease Risk Factors: |  |  |  |  |  |  |  |
| HDL Cholesterol | GLGC | Willer (2013) | 187167 | NA | NA | Mixed | SD (mg/dL) |
| LDL Cholesterol | GLGC | Willer (2013) | 173082 | NA | NA | Mixed | SD (mg/dL) |
| Total Cholesterol | GLGC | Willer (2013) | 187365 | NA | NA | Mixed | SD (mg/dL) |
| Triglycerides | GLGC | Willer (2013) | 177861 | NA | NA | Mixed | SD (mg/dL) |
| Cardiovascular Disease: |  |  |  |  |  |  |  |
| Coronary Heart Disease | CARDIoGRAMplus4CD | Nikpay (2015) | 184305 | 60801 | 123504 | Mixed | Binary |
| Myocardial Infarction | CARDIoGRAMplus4CD | Nikpay (2015) | 171875 | 43676 | 128199 | Mixed | Binary |
| Ischemic Stroke | ISGC | Malik (2016) | 21185 | 1859 | 19326 | European | Binary |
| Cardioembolic Stroke | ISGC | Malik (2016) | 21143 | 1817 | 19326 | European | Binary |
| Large Vessel Disease | ISGC | Malik (2016) | 20675 | 1349 | 19326 | European | Binary |
| Small Vessel Disease | ISGC | Malik (2016) | 29633 | 10307 | 19326 | European | Binary |
| **SECONDARY OUTCOMES:** |  |  |  |  |  |  |  |
| Circulating Metabolites | NA | Kettunen (2016) | 24925 | NA | NA | European | SD (mmol/L) |

**Abbreviations:** GLGC: Global Lipids Genetics Consortium; SSGAC: Social Science Genetic Association Consortium; UKB: UK Biobank; MRC-IEU: Medical Research Center-Integrative Epidemiology Center (UK Bristol); CARDIoGRAMplus4CD: Coronary ARtery Disease Genome wide Replication and Meta-analysis (CARDIoGRAM) plus The Coronary Artery Disease (C4D) Genetics; SD: standard deviation (units); mg/dL: milligram per decaliter; mmol/L: millimole per liter; HDL-C: high-density lipoprotein cholesterol; LDL-C: low-density lipoprotein cholesterol; SNP: single nucleotide polymorphism; CI: confidence interval.

**Fig 1.**
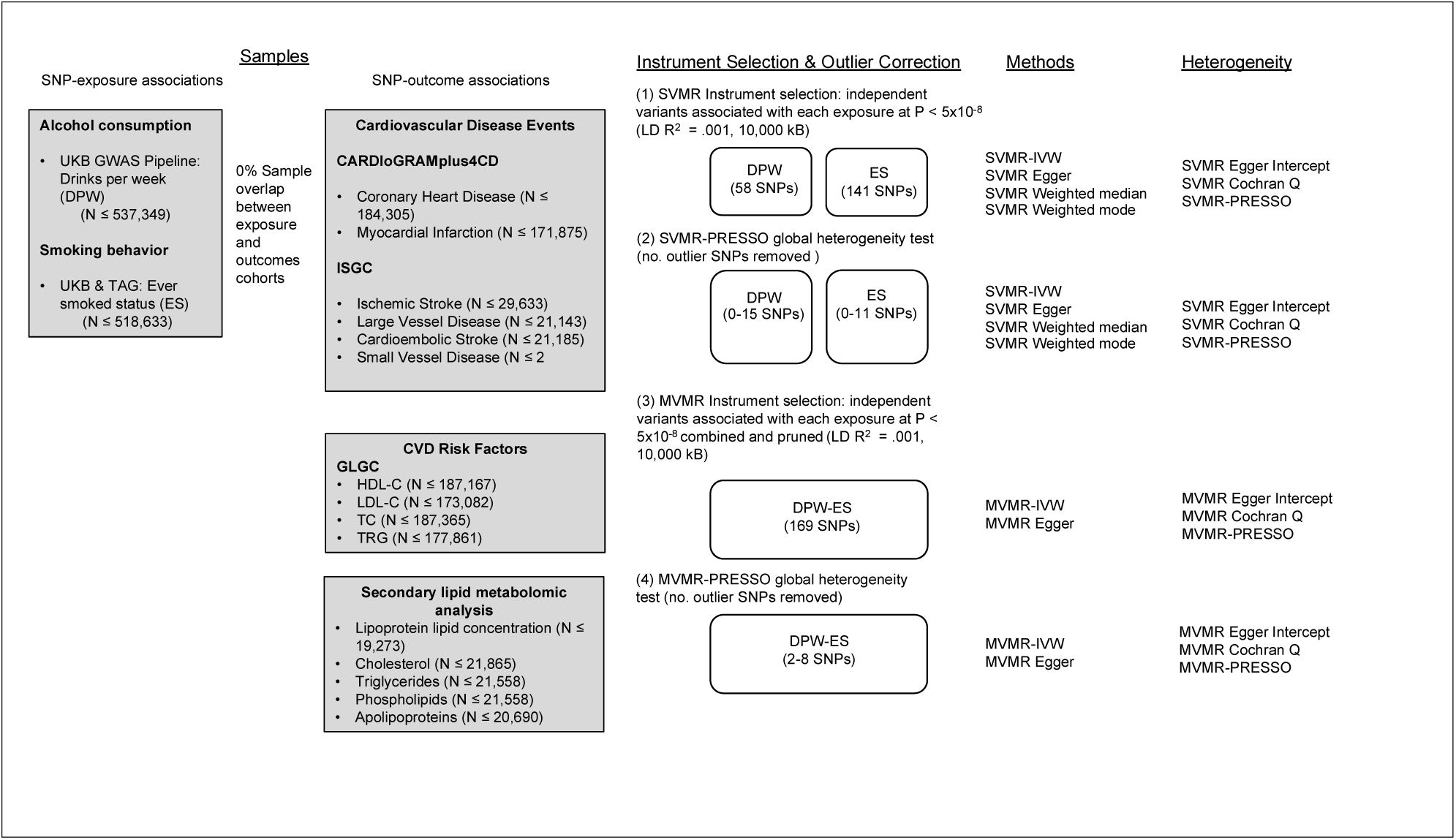
Study overview. Abbreviations: CVD; cardiovascular disease; Dx., diagnosis; GLGC: Global Lipids Genetics Consortium; SSGAC: Social Science Genetic Association Consortium; UKB: UK Biobank; MRC-IEU: Medical Research Center-Integrative Epidemiology Center (UK Bristol); CARDIoGRAMplus4CD: Coronary ARtery DIsease Genome wide Replication and Meta-analysis (CARDIoGRAM) plus The Coronary Artery Disease (C4D) Genetics; IVW, Inverse Variance Weighted MR; HDL-C: high-density lipoprotein cholesterol; LDL-C: low-density lipoprotein cholesterol; TC: total cholesterol; TRG: triglycerides; SNP: single nucleotide polymorphism. SVMR; single variable Mendelian randomization; MVMR; multivariable Mendelian randomization.

## RESULTS

### Effects of alcohol and tobacco use on cardiovascular disease lipid risk factors and events

In SVMR, we found genetic variants associated with increased alcohol use, not controlling for tobacco use, associated overall with increased levels of HDL-C (ß, 0.213, 95% CI, 0.101-0.325, *P*= 2.03×10^−4^), decreased levels of TRG (ß, -0.465, 95% CI, - 0.616- -0.314, *P*= 1.33×10^−9^), but not with levels of LDL-C nor TC. These results are shown in Fig. 2 and Table 2 in the Supplement. In MVMR, we found these variants, controlling for effects of tobacco use, associated with increased levels of HDL-C and decreased levels of TRG, but to a lesser extent (DPW: ß, 0.195, 95% CI, 0.053-0.091, *P*= 0.000; TRG: ß, -0.205, 95% CI, - 0.338, -0.072, *P*= 0.002). These results are shown in Fig. 2 and Table 3 in the Supplement. However, we did not find these variants associated, in either SVMR or MVMR, with risk of coronary heart disease (CHD), myocardial infarction (MI), nor any stroke, including ischemic, cardioembolic, large artery, and small vessel stroke.

**Fig 2.**
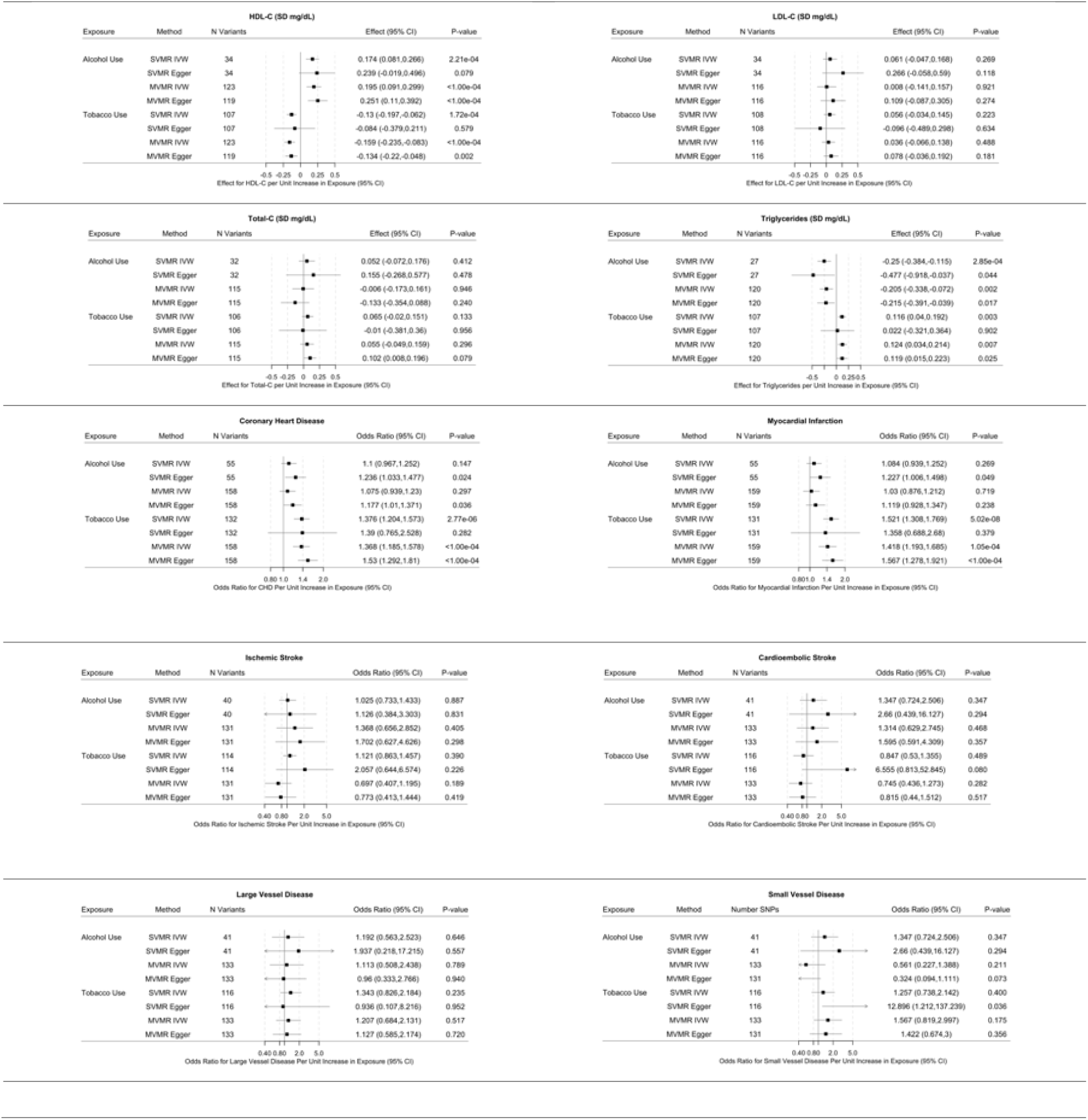
Multivariable Mendelian randomization results of alcohol and tobacco use on lipid cardiovascular disease risk factors and disease events. Potentially pleiotropic outlying SNPs removed. SVMR estimates total effect of a change in each of alcohol and tobacco use in 10 cardiovascular disease risk and event outcomes. MVMR estimates direct effect of alcohol and tobacco use conditional on tobacco and alcohol use respectively. MVMR inverse weighted variance methods shown with complementary MVMR Egger methods for SVMR extended to correct for both measured and unmeasured pleiotropy in MVMR. Full MR results presented in Tables 4, 5 and 6 in Supplement. Abbreviations: SVMR: single variable Mendelian randomization; MVMR: multivariable Mendelian randomization; HDL-C: high-density lipoprotein cholesterol; LDL-C: low-density lipoprotein cholesterol; SNP: single nucleotide polymorphism; CI: confidence interval; CVD: cardiovascular disease: IVW: inverse variance weighted MR.

In SVMR, we found genetic variants associated with increased risk of tobacco use (ever smoker status) associated, not controlling for alcohol use, with decreased levels of HDL-C (ß, -0.145, 95% CI, -0.218- -0.072, *P*= 1.15×10^−4^), increased levels of TRG (ß, 0.137, 95% CI, 0.057 -0.217, *P*= 8.31×10^−4^), but not with levels of LDL-C nor TC. We also found these variants associated with increased risk of CHD (OR, 1.309, 95% CI, 1.143-1.498, *P*= 1.05×10^−5^) and MI (OR, 1.48, 95% CI, 1.25-1.752, *P*= 4.88×10^−6^). We did not find these variants associated with any stroke, nor with ischemic, cardioembolic, and small vessel stroke. These results are shown in Fig. 2 and Table 4 in the Supplement. In MVMR, we found these variants, controlling for alcohol use, associated with decreased levels of HDL-C (ß, -0.159, 95% CI, -0.083- -0.235, *P*= 1.15×10^−4^), increased levels of TRG (ß, 0.124, 95% CI, 0.034 -0.214, *P*= 8.31×10^−4^), although again to a lesser extent than in SVMR. We found these variants, controlling for alcohol use, associated with increased risk of CHD (OR, 1.368, 95% CI, 1.185-1.578, *P*= 0.0000) and MI (OR, 1.48, 95% CI, 1.193-1.685, *P*= 1.05×10^−4^). These results are shown in Fig. 2 and Table 3 in the Supplement. We did not find these variants associated with any stroke, nor with ischemic, cardioembolic, and small vessel stroke, under either SVMR or MVMR.

MR-Egger can detect, under certain assumptions, directional pleiotropy; MR-Egger estimates are robust to such pleiotropy. While the magnitude and direction of the MR IVW effect estimates were not substantially different under SVMR and MVMR, the MR-Egger estimates were more precise under MVMR, simultaneously estimating the effects of alcohol and tobacco use. Further, the MR IVW effect estimates were more consistent (similar in magnitude and direction) with these MR-Egger estimates under MVMR analysis, strengthening our hand inferring causality from the MVMR results (Fig. 2 and Tables 2-4 in the Supplement).

### Effects of alcohol and tobacco use on lipoprotein and lipid metabolites

In exploratory MVMR analyses (reported here given our main interest in the direct versus the total (direct plus indirect) effects of alcohol and tobacco use), we found genetic variants associated with increased alcohol use, controlling for tobacco use, associated with increased concentrations of IDL particles (ß, 0.253, 95% CI, 0.065-0.441, *P*= 0.009), large LDL particles (ß, 0.280, 95% CI, 0.088-0.472, *P*= 0.004), medium LDL particles (ß, 0.253, 95% CI, 0.061-0.445, *P*= 0.012), small LDL particles (ß, 0.267, 95% CI, 0.069-0.4465, *P*= 0.008); very large HDL particles (ß, 0.285, 95% CI, 0.087-0.483, *P*= 0.002), large HDL particles (ß, 0.396, 95% CI, 0.218-0.574, *P*= 0.000), and medium HDL particles (ß, 0.248, 95% CI, 0.072-0.424, *P*= .011), but not with small HDL particles. These results are shown in Fig. 3, column 1, and Table 5 in the Supplement.

**Fig 3.**
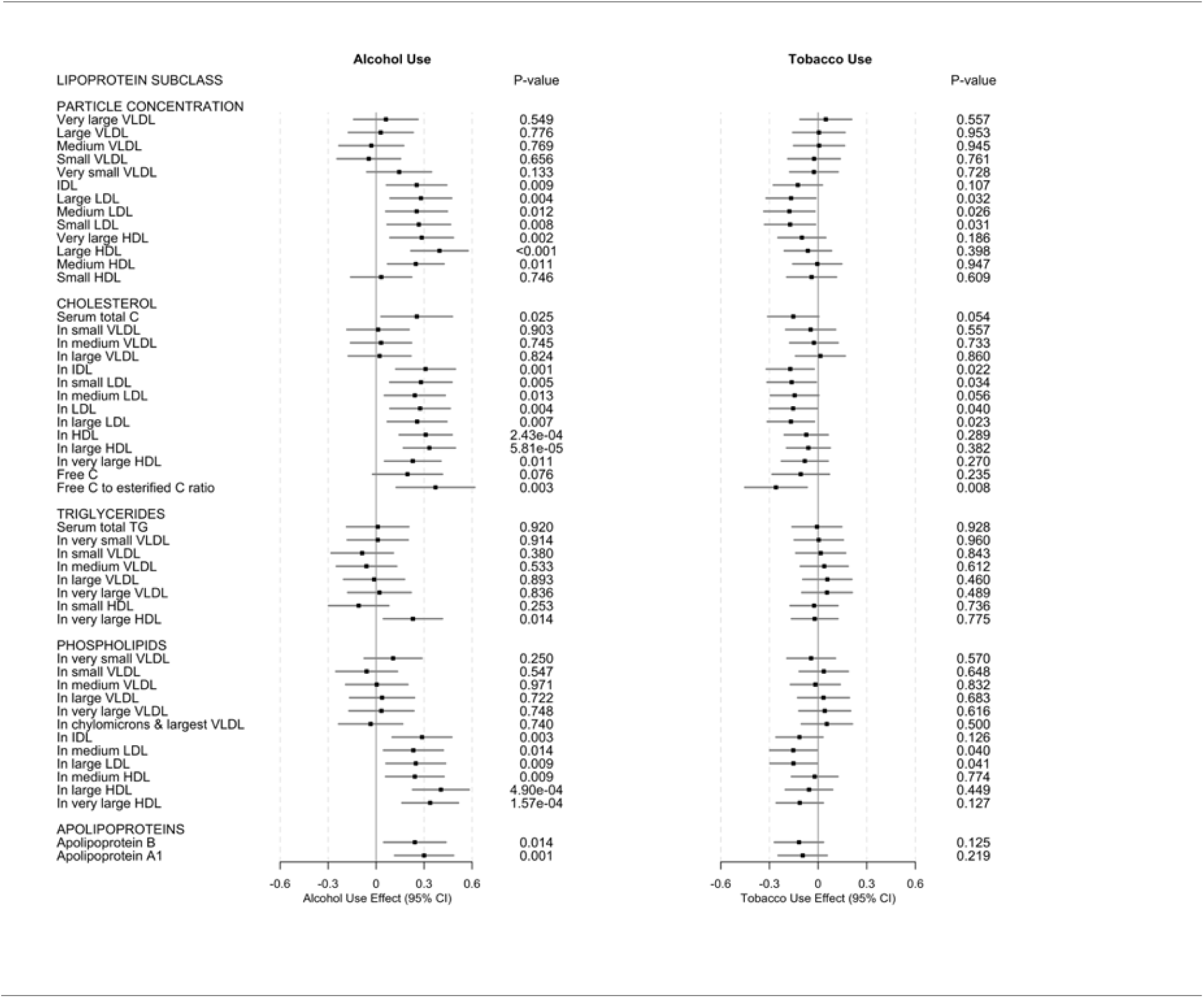
Multivariable inverse variance weighted Mendelian randomization (MVMR IVW) results for lipoprotein and lipid changes associated with alcohol and tobacco use. Results are shown as effects in SD-scaled concentration units per unit change in exposure alcohol or tobacco use. Full IVW MR results in Table 7 in the Supplement. Abbreviations: C: cholesterol; CI confidence interval; HDL: high-density lipoprotein; IDL: IDL: intermediate-density lipoprotein; LDL: low-density lipoprotein; PL: phospholipids; VLDL: very-low-density lipoprotein; TRG: triglycerides.

These alcohol use variants were further associated with increased levels of total cholesterol in IDL (ß, 0.309, 95% CI, 0.123-0.495, *P*= 0.001), small LDL (ß, 0.280, 95% CI, 0.086-0.474, *P*= 0.005), medium LDL (ß, 0.242, 95% CI, 0.052-0.432, *P*= 0.013), LDL (ß, 0.275, 95% CI, 0.087-0.463, *P*= 0.004) and large LDL (ß, 0.256, 95% CI, 0.070-0.442, *P*= 0.007); also HDL (ß, 0.310, 95% CI, 0.145-0.475, *P*= 2.43×10^−4^), large HDL (ß, 0.333, 95% CI, 0.171-0.495, *P*= 5.81×10^−5^), and very large HDL (ß, 0.229, 95% CI, 0.053-0.405, *P*= 0.011); not with levels of free cholesterol, but with free cholesterol to esterified cholesterol ratio (ß, 0.371, 95% CI, 0.126-0.616, *P*= 0.003). We did not find these variants associated with any triglyceride metabolites, except for triglycerides in very large HDL ((ß, 0.230, 95% CI, 0.046-0.414, *P*= 0.014). These results are shown in Fig. 3, column 1, and Table 5 in the Supplement.

Further still, these alcohol use variants were associated with increased levels of phospholipids in IDL (ß, 0.277, 95% CI, 0.101-0.473, *P*= 0.003); medium LDL (ß, 0.233, 95% CI, 0.047-0.419, *P*= 0.014); large LDL (ß, 0.248, 95% CI, 0.062-0.434, *P*= 0.009); and medium HDL (ß, 0.242, 95% CI,0.060-0.424, *P*= 0.009), large HDL (ß, 0.405, 95% CI, 0.229-0.581, *P*= 4.90×10^−6^), and very large HDL (ß, 0.338, 95% CI, 0.162-0.514, *P*= 4.90×10^−6^); as well as increased levels of apolipoprotein B (ß, 0.242, 95% CI, 0.048-0.436, *P*= 0.14) and A-1 (ß, 0.300, 95% CI, 0.116-0.484, *P*= 0.001). These results are shown in Fig. 3, column 1, and Table 5 in the Supplement.

Genetic variants associated with risk of tobacco use, controlling for alcohol use, were associated with fewer metabolites than, and in the direction opposite the alcohol use variants: We found tobacco use variants associated with decreased concentrations of large LDL particles (ß, -0.167, 95% CI, -0.320--0.014, *P*= 0.032), medium LDL particles (ß, -0.177, 95% CI, -0.334--0.020, *P*= 0.026), and small LDL particles (ß, -0.173, 95% CI, -0.330--0.016, *P*= 0.031); cholesterol in IDL (ß, -0.172, 95% CI, -0.319--0.025, *P*= 0.022), small LDL (ß, -0.163, 95% CI, -0.314--0.012, *P*= 0.040); medium LDL (ß, - 0.145, 95% CI, -0.294--0.004, *P*= 0.056), LDL (ß, -0.154, 95% CI, -0.301--0.007, *P*= 0.026), and large LDL (ß, -0.168, 95% CI, -0.313--0.023, *P*= 0.023), and also free cholesterol to esterified cholesterol ratio (ß, -0.260, 95% CI, -0.452--0.068, *P*= 0.008). We also found tobacco use variants associated with decreased levels of phospholipids, but only in medium LDL (ß, -0.153, 95% CI, -0.298--0.008, *P*= 0.040) and large LDL (ß, -0.152, 95% CI, -0.297--0.007, *P*= 0.041), but not in any triglycerides or apolipoproteins. These results are shown in Fig. 3, column 2, and Table 5 in the Supplement.

## DISCUSSION

Our two-sample MVMR design allowed us to estimate the direct effects of alcohol and tobacco use on cardiovascular disease risk factors and certain risk events, as opposed to and distinguished from the total effects estimated by single variable design. Specifically, total alcohol use was associated with increased levels of HDL-C, but decreased levels of triglycerides. Extending traditional lipid measures, our study also provides a comprehensive examination of the associations of alcohol and tobacco use on lipid particles and lipid constituents. In general, our findings support previous observational research showing a relationship between alcohol, smoking, and CVDs ^5,13^, and the consistent estimates across complementary MR methods and after controlling alcohol consumption for smoking (and smoking for alcohol consumption) strengthen causal inference. Finding robust increases in HDL-C with no corresponding reduction in CVD risk calls into question the mechanism through which observational literature attributed any cardioprotective effect of light-to-moderate alcohol consumption^14^. Similarly, our results, when combined with traditional RCTs and MR studies that failed to find CVD risk reduction of increased HDL-C ^34-36^ suggests that there is no cardioprotective effect of alcohol consumption.

Importantly, in our main analysis we failed to find a cardioprotective effect of alcohol use on CVD outcomes, suggesting that previously reported reductions in CVD risk due to other lifestyle differences that are associated with more light-to-moderate drinking patterns, such as healthier lifestyles^37^, are contributing to these effects. Our total weekly consumption instrument may have failed to differentiate between light-to-moderate drinking and heavier consumption, which are reported to have inverse associations with CVD disease risk in observational literature^14^. More likely, however, previous research may not have accounted for smoking, which we found to be significantly associated with increased CVD risk. For example, we failed to replicate an early MR study that found carriers of the *ADH1B* rs1229984 A-allele (associated with reduced alcohol consumption) were also associated with lower CHD and stroke risk^8^. However, they reported that rs1229984 was also associated with smoking: omitting smoking from the alcohol consumption equation would have influenced their results. They also found rs1229984 was associated with increased educational attainment, which more recent MR studies have shown to be causally linked with CVD risk^38^. Expanding upon their single stroke variable, which may mask differential associations between alcohol and stroke subtypes^11^ further strengthens our conclusions. Failing to replicate the instrumental variable estimates Millwood et al. 2019 found among participants in the China Kadoorie Biobank sample supports the hypothesized differential effect of alcohol consumption on CVD risk across ethnic backgrounds^14,39^. East Asia has the highest prevalence of stroke worldwide^40^, and generally lower alcohol consumption behaviors than European countrie.^3^. In addition, survey data show that men consume 95% of all alcohol in China^41^. Therefore, caution is needed before extrapolating these findings to other populations and our study highlight the need for population-specific MR analysis in elucidating the underlying alcohol-CVD relationship.

Consistent with results derived from observational literature, we found strong genetic evidence of a causal effect of tobacco use, independent of alcohol use, on increased MI and CHD risk^5^. Our estimates showing 48% and 30% increased risk for MI and CHD are similar to estimates from observational and smoking cessation studies ^42^, and we confirm findings from MR study that found a smoking instrument constructed from multiple smoking behaviors increased CHD and MI risk^38^ by replicating their results using a separate smoking instrument accounting for alcohol consumption. Mechanisms underlying the association with MI may include nicotine-induced sympathetic nervous system activation and vascular endothelial function impairment, which together may induce cardiovascular stress through increased blood pressure, heart rate, and cardiac output^43^, while long-term effects on atherosclerotic plaques, thrombosis, and inflammatory processes may mediate CHD risk^5^. While an increased awareness and understanding of the adverse health effects associated with smoking has decreased the prevalence of current smoking, smoking continues to be a major cardiovascular hazard^43^. For example, in the United States, while the percentage of adults reporting having ever smoked has decreased from 43% in 1965 to 19% in 2010^43^, 44 million adults still smoke. We were unable to assess the causal effect of smoking on CVD mortality; however, a meta-analysis found smoking cessation associated with a 36% reduced mortality risk, which is similar in magnitude to other prevention strategies, including statins^44^, and ß-blockers^44^. If the association is causal, our results may be useful to further inform continued clinical trial and smoking prevention program development.

We failed to replicate the increase in stroke risk associated with smoking Carter et al. 2019 found using a constructed smoking instrument among UK Biobank and MEGASTROKE participants^38^. Our smoking instrument was based upon the “ever smoker status”, selected to pick up generalized lifetime smoking behavior for our primary alcohol use analysis; to the extent the “smoker” status included current “non-smokers”, we would expect some impact on the effect of smoking on risk of stroke risk. Given the strong evidence of a substantial adverse effect of even one cigarette per day on risk of CVD and stroke^45^, caution is needed when interpreting our null finding.

Our findings also support the causal role of alcohol consumption in increasing HDL-C as reported in previous MR studies^18,26^ and provides evidence for the finding after controlling for smoking behavior. While we were unable to evaluate the effects of heavy alcohol consumption on HDL-C levels, we recently reported a cross-sectional study findings that very heavy alcohol consumption (> 10 standard drinks for men and > 8 standard drinks for women) to be associated with increased HDL-C^46^, which suggests that alcohol may impact HDL-C levels across a range of alcohol consumption levels. Future MR studies are needed to investigate this hypothesis. Our comprehensive examination of the associations of alcohol with lipid particles and lipid constituents replicated a recent metabolomic study^7^, and revealed that alcohol may increase large HDL-C particle size, total cholesterol concentration and apolipoprotein A1, which each are associated with reduced MI risk^15^; however, when considering coincident increases in MI increasing constituents such as small and medium LDL-C and large LDL-C particle size^15^, and, as mentioned above, the RCTs and MR studies failing to find increased HDL-C and reduced CVD risk^8,34-36^, further suggests any hypothetical cardioprotective mechanism derived from HDL-C metabolism changes may be further reduced when compared to assessments failing to consider these heterogenous associations. It is possible that alcohol consumption may reduce CHD risk through its modulation of HDL-C phospholipid content, which impacts reverse cholesterol transport^47^– a pathways inversely associated with prevalent CHD, even after controlling for HDL-C levels^48^ – and has been shown to be reduced among CHD patients^47^.

Our finding that alcohol use reduces TRG confirms results from MR studies using European^8,18^ and European American^26^ cohorts; however, our 58 SNP instrument for alcohol use may account, in part, for the reported direction. The association between alcohol and triglycerides remains equivocal^14^ with a meta-analysis of intervention studies reporting no association^13^, and observational studies have reported alcohol associated with either reduced, or harmful TRG levels, depending on the cohort and the amount consumed^49^. However, the intervention studies investigated primarily short-term effects while genetic variants associated with alcohol consumption are constant throughout the life course^18^. Since our finding was also in a European American sample, and alcohol may affect TRG levels differently depending on race or ethnicity^50^, future trans-ancestry replication studies are needed.

Supporting the observational literature^5^, we found genetic evidence for a causal effect of smoking on decreased levels of HDL-C and increased levels of TRG. Conversely, our findings do not support Åsvold et al. 2014’s MR analysis of the Norwegian HUNT Study cohort showing higher HDL-C and no effect on triglycerides using a single SNP instrument (rs1051730 located in *CHRNA3*)^51^. However, associations between rs1051730 and HDL-C were not significantly different by smoking status in the HUNT cohort, and even the authors cautioned interpretation of their HDL-C finding^51^. Smoking has been shown to alter lipid metabolism enzymes, which would alter HDL-C levels^52^. Further, HDL-C particles can be rendered nonfunctional due to oxidative modifications from smoking, which suggests smoking may impact HDL-C quantity and function^52^.

Our study has several strengths. We use a multivariable two-sample MR design, the most appropriate design given the strong correlation between alcohol and tobacco use, yielding unbiased and consistent estimates of the direct effects of our multiple exposures on the CVD risk factors and disease events^24^. We selected the SNPs included in the genetic instruments through a stringent process. Another strength is the use of large publicly available summary-level genetic study datasets (from more than 940,000 study participants overall) that are necessary for two-sample MR^53^ and the broadly consistent estimates across multiple MR methods, each relying on orthogonal assumptions, provides confidence in result robustness and strengthens causal inference^25^. We also found no evidence of horizontal pleiotropy, suggesting minimal potential bias^22^. Finally, while previous MR studies only used a combined stroke variable, we used multiple stroke subtypes, given some observational studies reporting a differential association of alcohol by stroke subtype^11^, which would strengthen our conclusion regarding a null effect of alcohol use on stroke risk. Similarly, our metabolomic analysis highlighting the complex impact of alcohol use on circulating metabolites may be an important addition to the field’s growing body of MR literature.

Several limitations should be noted. As with previous alcohol literature, alcohol consumption patterns are variable, and the exposure variables may be either under- or over-reported^27^. Carter et al. 2019 found that smoking mediates the relationship between education and CVD risk^38^, and we recently reported that a two-sample MR finding educational attainment (EA) affects how alcohol is consumed – with increased EA increasing whether alcohol is consumed with meals, drinking more frequently, but less likely to binge drink (≥6 drinks per occasion)^54^. Considering previous work has shown that how alcohol is consumed is associated with different dietary patterns^55^, and differences in CVD risk^28^, it is possible dietary patterns may moderate the association between alcohol and CVD risk. Therefore, future studies using cohorts with heavier reported consumption or incorporating dietary patterns associated with different drinking typologies would further elucidate the alcohol-CVD relationship.

While genetic instruments are considered to represent lifelong exposure^22^, currently no GWAS available examines the genetic architecture of alcohol consumption throughout the entire life course. In addition, while we show an effect of alcohol consumption and smoking on certain CVD risk factors, alcohol has been associated with other biological mechanisms, such as, such as cellular adhesion molecules and adipocyte hormones, that may contribute to CVD risk^13,14^, which we did not assess. We examined total alcohol consumption among participants in the UK Biobank, who have been reported to be more educated, have healthier lifestyles, and fewer health problems than the UK population^56^, and other hazardous consumption patterns associated with increased CVD risk^57^, were not assessed..

Further, the included GWAS are from survey participants of European ancestry, and given observational evidence suggesting different associations between alcohol and CVDs exist across racial and ethnic groups^14^, caution may be advised in extrapolating these findings to other populations. Genome-wide significant (GWS) SNPs for each exposure only explain a small fraction of total variance in complex traits^58^. Relatedly, some GWS exposure SNPs, or their proxies, were not available in every outcome GWAS used in this study, suggesting reduced variance captured with the remaining SNPs^25^. Also, given the absence of available outcome GWASs, we were unable to replicate the findings in independent datasets. While we found no evidence of pleiotropy, it is possible pleiotropy still exists from unmeasured confounders. Relatedly, while unlikely, we cannot entirely rule out the possibility of sample overlap between exposure and outcome datasets, which would bias estimates^59^; however, any bias would likely be minimal^59^.

Using two-sample multivariable MR, we show a likely direct causal effect of alcohol consumption on increased HDL-C, and we failed to find a cardioprotective effect of alcohol. However, smoking significantly increased the risk for MI, CHD, and stroke. Our findings suggest previous observational studies reporting an association between alcohol consumption and CVDs may have resulted from confounding, and differences between our findings and recent MR analyses using data from populations with different ethnic background suggest that future MR studies in additional populations may help elucidate this complex relationship. Our exploratory metabolomic analyses highlight the complex impact of alcohol on both cardioprotective biomarkers and biomarkers associated with increased CVD risk. Our findings may help inform the public health debate regarding the alcohol-CVD relationship and suggest that future work examining this relationship must account for smoking behaviors. In conjunction with other study designs, this study provides additional evidence that programs targeted at reducing alcohol consumption and smoking may have beneficial cardiovascular effects.

## METHODS

### Data sources

We conducted single-variable (SV) as well as multivariable (MV) two-sample MR analyses of alcohol and tobacco use on cardiovascular disease and disease risk factors using summary-level data from publicly available GWASs from similar populations of predominantly European ancestry of each of alcohol and tobacco use and cardiovascular disease and disease risk factors (Table 1 and Table 1 in the Supplement). All studies have existing ethical permissions from their respective institutional review boards and include participant informed consent and included rigorous quality control (Methods 1 in the Supplement). Figure 1 summarizes the study design.

### Instruments

SNPs for alcohol use were extracted from a large GWAS meta-analysis of 414,343 individuals of European ancestry from the UK Biobank prospective cohort study collected across the United Kingdom from 2006-2010. We included all SNPs associated at genome-wide significance (*P* < 5 × 10^−8^) (we use the standard heuristic significance threshold for these dimensional MR to make the work manageable) with the phenotype “drinks per week” (DPW), constructed, for UK Biobank participants who indicated they drank “at least once or twice per week”, by aggregating the weekly intake of distilled spirts (pub measures), beer and cider (pints), red wine, white wine, and champagne (glasses), and other alcoholic drinks *e*.*g*. alcopops (DPW: mean = 8.92 drinks, SD = 9.30 drinks)^32^. (For UK Biobank participants who indicated they drank “one to three times a month”, the phenotype was constructed by aggregating the monthly intake over all drink types and dividing by four.) We pruned the results to exclude all SNPs with a pairwise linkage disequilibrium (LD) R^2^ > 0.001, giving us 58 independent SNPs associated with alcohol use for the single-variable MR (SVMR) analyses (Table 6 in the Supplement).

For tobacco use, we extracted SNPs from the companion GWAS meta-analysis of 518,633 individuals of predominantly European ancestry (444,598 from the UK Biobank prospective cohort study and 74 035 from the Tobacco, Alcohol and Genetics (TAG) Consortium)^32^. We included all SNPs associated at genome-wide significance with the phenotype “ever smoker status” (a binary phenotype coded “1” if the individual reported he or she was a previous or current smoker, “0” if he or she had never smoked or only smoked once or twice). We pruned the results to exclude all SNPs with a pairwise LD R^2^>.001, giving us 141 independent SNPS associated with tobacco use for SVMR analyses (Table 7 in the Supplement).

For the multivariable MR (MVMR) analyses, we included all 199 SNPs which were genome-wide significant in either the GWASs of either alcohol or tobacco use, deleting 2 duplicate SNPs (SNPs associated at genome-wide significance with both alcohol and tobacco use). (We note as regards number of unique instruments for each exposure, that 14 of the 58 instruments for alcohol use were associated with tobacco use (at nominal P of 0.05 corrected for 58 comparisons); 16 of the 141 instruments for tobacco use were associated with alcohol use (at nominal P of 0.05 corrected for 141 comparisons)). We pruned the results to exclude any SNPs with a pairwise LD R^2^ > 0.001, giving us 169 independent SNPs for the MVMR analyses.

### Cardiovascular risk factors

We used summary statistics from the GWAS meta-analyses for lipid levels – HDL-C, LDL-C, TC and triglycerides (measured in SD units (mg/dL)) – in 37 cohorts (none UKB nor TAG) comprising 188,577 persons of predominantly European ancestry (excluding persons known to be on lipid lowering medications or women known to be pregnant)^60^. Of the 58 possible genome-wide significant SNPs associated with alcohol use, 28 SNPs were present in the GWASs, 30 SNPs were missing, 14 SNPs were identified in high linkage disequilibrium (LD) as proxies for some of the missing SNPs were present, 5 SNPs were removed during harmonization for being palindromic with intermediate allele frequency, leaving 37 SNPs for SVMR analysis. Of the 141 possible SNPs associated with tobacco use, 98 SNPs were present in the GWASs, 43were missing, 23 SNPs were identified in high linkage disequilibrium (LD) as proxies for some of the missing SNPs were present, and 11 SNPs were removed during harmonization for being palindromic with intermediate allele frequency, leaving 110 SNPs for SVMR analysis. Of the 169 possible SNPs of the combined instrument set, 110 SNPs were present in the GWASs, 59 were missing, 27 SNPs were identified in high linkage disequilibrium (LD) as proxies for some of the missing SNPs were present, and 14 SNPs were removed during harmonization for being palindromic with intermediate allele frequency, leaving 123 SNPs for MVMR analysis.

### Cardiovascular disease events

We used summary statistics from the GWAS meta-analyses for coronary artery disease (CAD: 60,801 cases; 123,504 controls) and myocardial infarction (CAD sub-group: 43 676 cases; controls 128,199 controls) in 48 cohorts (none UKB nor TAG) in up to 184 305 persons of mixed ancestry (approximately 15% non-European ancestry, including Chinese, Indian Asian, South Asian, Lebanese, African-American and Hispanic American ancestry)^61^. Of the 58 possible SNPs associated with alcohol use, all SNPs were present in the GWASs, but 1 was removed during harmonization as palindromic, leaving 57 for SVMR analysis. Of the 141 possible SNPs associated with tobacco use, 140 SNPs were present in the GWASs, and 6 SNPs were removed during harmonization as palindromic with intermediate allele frequency, leaving 134 SNPs for SVMR analysis. Of the 169 possible SNPs of the combined instrument set, 167 SNPs were present in the GWASs, 2 were missing, 1 SNP was identified in high linkage disequilibrium (LD) as proxies for some of the missing SNPs were present, and 6 SNPs were removed during harmonization for being palindromic with intermediate allele frequency, leaving 162 SNPs for MVMR analysis.

For ischemic stroke and ischemic stroke sub-types, we used summary statistics from the GWAS meta-analyses in 12 cohorts in up to 29,633 persons of European ancestry^62,63^. Of the 58 possible SNPs associated with alcohol use, 28 SNPs were present in the GWASs, 14 SNPs were identified in high linkage disequilibrium (LD) as proxies for some of the missing SNPs, and 1 SNP was removed during harmonization for being palindromic with intermediate allele frequency, leaving 41 SNPs for SVMR analysis. Of the 141 possible SNPs associated with tobacco use (ever smoker status), 100 SNPs were present in the GWASs, 20 SNPs were identified in high linkage disequilibrium (LD) as proxies for some of the missing SNPs, and 5 SNPs were removed during harmonization as palindromic with intermediate allele frequency, leaving a total of 116 SNPs for analysis. Of the 169 possible SNPs of the combined instrument set, 112 SNPs were present in the GWASs, 57 were missing, 26 SNPs were identified in high linkage disequilibrium (LD) as proxies for some of the missing SNPs were present, and 5 SNPs were removed during harmonization for being palindromic with intermediate allele frequency, leaving 133 SNPs for MVMR analysis.

### Circulating metabolites

We used summary statistics from the GWAS meta-analyses of 14 cohorts (none UKB nor TAG) in up to 24,925 persons of European ancestry for 58 phenotypes describing circulating metabolites (under 5 lipoprotein sub-groups headings *i*.*e*. particle concentrations, triglycerides, cholesterol, phospholipids and apolipoproteins (see Table 1 in the Supplement) ^64^. Of the 169 possible SNPs of the combined instrument set for exploratory MVMR, 162 SNPs were present in the GWASs, 7 were missing, 6 SNPs were identified in high linkage disequilibrium (LD) as proxies for some of the missing SNPs were present, and 6 SNPs were removed during harmonization for being palindromic with intermediate allele frequency, leaving 162 SNPs for MVMR analysis.

### Sample overlap

Participant overlap in samples used to estimate genetic associations between exposure and outcome in two sample MR can bias results^59^. We endeavored to use only non-overlapping GWAS summary statistics to reduce this source of weak instrument bias; we avoided overlap between the primary exposure and outcomes cohorts included.

### Statistical analysis

For SVMR, we used inverse-variance weighted MR (MR IVW) (single variable weighted linear regression) along with the complementary MR-Egger method to assess the evidence of the causal effects of each of alcohol and tobacco use on cardiovascular disease risk factors and disease outcomes and to detect the sensitivity of the results to different patterns of violations of IV assumptions^22^: consistency of results across methods strengthens an inference of causality^22^. For MVMR, introduced by Burgess et al.^23^, we used an extension of the inverse-variance weighted MR method, performing multivariable weighted linear regression (variants uncorrelated, random-effect model) with the intercept term set to zero^65^. We also used an extension of the MR-Egger method to correct for both measured and unmeasured pleiotropy^66^.

### Sensitivity analyses

To evaluate heterogeneity in instrument effects, which may indicate potential violations of the IV assumptions underlying two-sample MR^67^, we used the MR Egger intercept test^67^, and the Cochran heterogeneity test^68^, and multivariable extensions thereof^65,66^. The MR <u>p</u>leiotropy <u>re</u>sidual <u>s</u>um and <u>o</u>utlier (MR-PRESSO) global test, and multivariable extension thereof^69^, were used to facilitate identification and removal of outlier variants in order to correct potential directional horizontal pleiotropy and resolve detected heterogeneity. Details about MR methods and tests used are included in Methods 2 in the Supplement. For the SVMR, we used the Steiger directionality test to test the causal direction between the hypothesized exposure and outcomes^70^. Given a nominal threshold of .05, with 11 comparisons in the primary analysis, the Bonferroni corrected threshold for the study would be .00454. Analyses were carried out using MendelianRandomization, version 0.4.1, TwoSampleMR, version 4.16^22^, and MR-PRESSO, version 1.0^69^, in the R environment, version 3.5.1.

### Reported results

We generally look for those estimates (1) substantially agreeing in direction and magnitude across MR methods, (2) exceeding nominal significance corrected for ten (10) multiple comparison (*P* < 0.005) in MR IVW, (3) not indicating bias from horizontal pleiotropy (MR-PRESSO global *P* > 0.05, also MR Egger intercept *P* > 0.05), and for SVMR (4) indicating true causal effect directionality (Steiger directionality test *P* < 0.05). Full MR results with test statistics both before and after outlier correction are presented in Tables 2-5 in the Supplement. MR-PRESSO outlier corrected results are displayed in Figures 2 and 3. Causal effects estimates are presented as the MR IVW (with MR-Egger as the complementary method presented to draw inferences about causality) effect estimate (ß) (with 95% confidence interval (CI)), or, for the binary cardiovascular artery disease outcomes, as the odds ratios (OR) (with 95% CI), per unit increase in the exposures, *i*.*e*. one-unit increase in drinks per week (DPW); or a one-unit increase in the logarithm of the odds ratio (log(OR) of the risk of ever smoking tobacco (ever smoker status).

## ACKNOWLEGEMENTS

This research was facilitated by the Social Science Genetic Association Consortium (SSGAC), the Substance Use Disorders Working Group of the Psychiatric Genomics Consortium (PGC-SUD) (supported by funds from NIDA and NIMH to MH109532 and, previously, with analyst support from NIAAA to U01AA008401 (COGA)), and the Medical Research Council Integrative Epidemiology Unit (MRC-IEU, University of Bristol, UK), especially the developers of the MRC-IEU UK Biobank GWAS Pipeline. We gratefully acknowledge their contributing studies and the participants in those studies without whom this effort would not be possible. This work was supported by the National Institutes of Health (NIH) intramural funding [ZIA-AA000242 to F.W.L]; Division of Intramural Clinical and Biological Research of the National Institute on Alcohol Abuse and Alcoholism (NIAAA). The authors declare no conflict of interest.

## DATA AVAILABILITY

All analyses were conducted using publicly available data. The summary-level data for alcohol and tobacco use are available at https://www.thessgac.org/data, and the summary-level data for lipids (GLGC), coronary heart disease (CARDIoGRAMplus4CD), stroke (ISGC), and metabolites are all available through MR Base at http://www.mrbase.org/

## CODE AVAILABILITY

The analysis code in R is available on request and all data displayed in the figures are available in the Supplementary Tables.

## TABLES & FIGURES

**Table 1.** Phenotypes sources

**Fig 1**. Study overview

**Fig 2**. Multivariable Mendelian randomization results of alcohol and tobacco use on lipid cardiovascular disease risk factors and disease events

**Fig 3**. Multivariable inverse variance weighted Mendelian randomization (MVMR IVW) results for lipoprotein and lipid changes associated with alcohol and tobacco use

