## Supplemental Methods for "Evaluating the effects of alcohol and tobacco use on cardiovascular disease using multivariable Mendelian randomization"

**Supplementary Information**

**Supplementary Methods** **1.** Data sources

**Supplementary Methods 2.** Statistical analysis

**Supplementary Table 1.** Phenotype source and description

**Supplementary Table 2.** Single variable Mendelian randomization results of SSGAC drinks per week in UKB participants on cardiovascular disease risk factors and cardiovascular disease

**Supplementary Table 3.** Multivariable Mendelian randomization results of SSGAC drinks per week and ever smoker status in UKB participants on cardiovascular disease risk factors and cardiovascular disease

**Supplementary Table 4**. Single variable Mendelian randomization results of SSGAC ever smoker status in UKB participants on cardiovascular disease risk factors and cardiovascular disease

**Supplementary Table 5.** Single- and multivariable inverse variance weighted Mendelian randomization results of alcohol consumption (DPW) and ever smoker status (ES) on phospholipids phenotypes

**Supplementary Table 6.** Instruments for exposure drinks per week aligned with ever smoker status

**Supplementary Table 7**. Instruments for ever smoker status aligned with drinks per week

### SUPPLEMENTARY METHODS

### 1. DATA SOURCES

**Alcohol consumption (Drinks per week) in the UKB: SSGAC UKB*.*** Full details of the methods applied for phenotype characterization, genotyping and imputation, quality control, and GWAS methods can be found in the SSGAC GWAS^1^. All association statistics used herein are from the meta-analysis of the drinks per week construct in the UKB (sample size N = 413,343). Only participants satisfying the following criteria were eligible for inclusion: (1) the participant was of European ancestry; (2) the participant passed the standard quality controls described in the Supplementary Materials accompanying publication of the GWAS; and (3) all relevant covariates were available for the participant. The GWAS was limited to 22 autosomes. Association analyses were conducted using linear regression; covariates included a vector of principal components of the genetic relatedness matrix after application of the pre-imputation filters described above; a vector of other control variables, including controls for sex and birth year, and sex-specific birth year fixed effects; and a vector containing cohort-specific controls and technical covariates (such as dummy variables for genotyping array and genotyping batches).

**Tobacco use (Ever Smoker Status) in the UKB: SSGAC UKB*.*** Full details of the methods applied for phenotype characterization, genotyping and imputation, quality control, and GWAS methods can be found in the SSGAC GWAS^1^. All association statistics used herein are from the meta-analysis of the ever smoker tobacco status (whether one has ever been a smoker) from the SSGAC GWAS on cohorts from UKB (N=444,598) and the Tobacco, Alcohol and Genetics (TAG) Consortium29 (N=74,035).^1^ Assessed as a binary phenotype, “ever smoker” was coded “1” if the sample participant reported they were a previous or current smoke, “0” if they had never smoked or only smoked once or twice. Only participants satisfying the following criteria were eligible for inclusion: (1) the participant was of European ancestry; (2) the participant passed the standard quality controls described in the Supplementary Materials accompanying publication of the GWAS; and (3) all relevant covariates were available for the participant. The GWAS was limited to 22 autosomes. Association analyses were conducted using linear regression; covariates included a vector of principal components of the genetic relatedness matrix after application of the pre-imputation filters described above; a vector of other control variables, including controls for sex and birth year, and sex-specific birth year fixed effects; and a vector containing cohort-specific controls and technical covariates (such as dummy variables for genotyping array and genotyping batches).

**Blood lipid levels: GLGC*.*** Full details of the methods applied for phenotype characterization, genotyping and imputation, quality control, and GWAS methods can be found in the GLGC GWAS^2^. All association statistics used herein are from the meta-analysis on blood-lipid levels in 188,577 participants from 60 studies, with potential participants known to be pregnant or on lipid-lowering medications excluded. Association analyses were conducted using linear regression to evaluate the additive effects of each SNP on blood lipid levels after adjusting for age and sex. Genomic control values for the initial meta-analyses were low for a sample of this size, indicating that population stratification would have a minor impact on results. After genomic control correction, 157 loci associated with blood lipid levels were identified (P < 5×10^−8^), several of which loci were validated by a similar extension based on prior GLGC GWAS results.

**Cardiovascular artery disease: CARDIoGRAMplus4CD*.*** Full details of the methods applied for phenotype characterization, genotyping and imputation, quality control, and GWAS methods can be found in the CARDIoGRAMplus4CD GWAS^3^. All association statistics used herein are from GWAS meta-analysis of CAD in 60,801 cases and 123,504 controls (N = 184,305) from 48 studies, with up 6.7 million common (MAF>0.05) as well as 2.7 million low frequency (0.005<MAF<0.05) variants. Three models of heritable disease susceptibility were analyzed by logistic regression; minor and major alleles were identified by reference to the allele frequencies in the pooled populations (*i.e.* all continents) of 1000 Genomes phase 1 v3 data. Variants that were retained in at least 60% of the studies were submitted for inverse-variance weighted fixed effects meta-analysis.

**Ischemic stroke and stroke sub-types: ISGC**. Full details of the methods applied for phenotype characterization, genotyping and imputation, quality control, and GWAS methods can be found in ISGC 2016 GWAS^4^. All association statistics used herein are from the discovery stage GWAS meta-analysis of stroke and 3 stroke sub-types in 10,307 Caucasian cases and 19,326 Caucasian population- matched controls from 12. Approximately 9 million SNPs and 1 million indels were available for analysis after QC. Promising signals were replicated both in Caucasian and non-Caucasian populations. Logistic regression was performed independently in each of the 12 samples; meta-analysis was performed centrally for all datasets. Covariates were not considered as they were not equally available over all studies.

**Circulating metabolites*.*** Full details of the methods applied for phenotype characterization, genotyping and imputation, quality control, and GWAS methods can be found in the GWAS^5^. All association statistics used herein are from the meta-analysis on 123 human blood lipid and circulating metabolite concentrations (only 49 analyzed in this study) quantified by high-throughput NMR spectroscopy metabolomics for up to 24 925 participants of European ancestry (Finnish, Estonian, Dutch and German) from 14 genotyped data sets (derived from 10 case-control, population-based, family-based, birth cohort, and follow-up in children studies): COROGENE, Genetic Predisposition of Coronary Heart Disease in Patients Verified with Coronary Angiogram; EGCUT, Estonian Genome Center of University of Tartu Cohort; ERF, Erasmus Rucphen Family Study; FR97, a subsample of FINRISK 1997; FTC, Finnish Twin Cohort; GenMets, Genetics of METabolic Syndrome; HBCS, Helsinki Birth Cohort Study; KORA, Cooperative Health Research in the Region of Augsburg; LLS, Leiden Longevity Study; *N*, number of individuals with both genotype and metabolite traits analysed; NFBC 1966, Northern Finland Birth Cohort 1966; NTR, Netherlands Twin Register; PredictCVD, FINRISK subsample of incident cardiovascular cases and controls; PROTE, EGCUT sub-cohort; YFS, The Cardiovascular Risk in Young Finns Study. Across the 14 cohorts, percent female ranged from 37% to 64%; age ranged from mean (standard deviation) 23.9 (2.9) to 61.3 (2.9) years. Cohort-specific adjustments for the meta-analysis included family structure, familial relations, relatedness, and genotyping sample. Sample quality control excluded sex mismatches, outlier by eye, relatedness, Mendelian inconsistencies, and heterozygosity. Cohorts were analyzed individually using linear mixed model association methods and summary statistics combined in a meta-analysis.

### 2. STATISTICAL ANALYSIS

Complementary methods including inverse variance-weighted MR (MR IVW), MR Egger, weighted median, and weighted mode MR were used to assess the causal effects of alcohol and tobacco use on cardiovascular disease risk factors and disease events and also circulating metabolites. MRIVW is generally regarded as the main method: in the absence of pleiotropy and assuming the instruments are valid, MRIVW estimates are the best unbiased estimates^6^. Consistency of results across methods (each making different assumptions about pleiotropy) strengthens causal inference; divergent results may indicate bias from genetic pleiotropy^6^.

**MR IVW**. For SVMR, implements a single weighted regression of the exposure SNP effects against the outcome SNP effects (meta-analyses SNP-specific Wald estimates from exposure and outcome GWASs), with the intercept term set to zero; the weights derived from the inverse outcome effects’ variance; multiplicative random effects, allowing SNPs to have different mean effects, allowing for heterogeneity due to e.g. horizontal pleiotropy; returns unbiased estimates of a causal effect are returned so long as horizontal pleiotropy is balanced^6,7^. For MVMR^8^, implements multivariable weighted linear regression; variants correlated or (as in this study) uncorrelated (pruning instruments to LD R^2^ < 0.001); intercept term set to zero^9^.

**MR Egger**. Extends MRIVW by not setting intercept to zero, allowing the net-horizontal pleiotropic effect across all SNPs to be unbalanced or directional (some SNPs could be acting on the outcome through a pathway other than through the exposure)^6,10^. MR Egger thus relaxes assumption of “no horizontal pleiotropy” and assumes only that horizontal pleiotropic effects are not correlated with SNP-exposure effects. MR Egger returns unbiased causal effect estimates even if assumption is violated for all SNPs, but estimates are less precise (wider confidence intervals expected) than MRIVW. MR Egger intercept estimates the directional pleiotropic effect ^6^.

**Weighted median MR**. Uses median effect of all available SNPs; only half the SNPs need to be valid instruments (no horizontal pleiotropy, no association with confounders, and robust association with the exposure) to return unbiased causal effect estimates. Stronger SNPs contribute more towards causal estimates, with the contribution of each SNP weighted by the inverse variance of its outcome association^11^.

**Weighted mode MR**. Clusters SNPs into groups based on similarity of causal effects; and returns the causal effect estimate based on cluster with largest number of SNPs; returns unbiased causal effects so long as SNPs within the largest cluster are valid instruments; weights each SNP’s contribution to the clustering by the inverse variance of its outcome effect. Assuming the most common causal effect is consistent, estimated causal effects would be unbiased even if all other instruments are invalid ^12^.

**Sensitivity analyses and diagnostics.** To evaluate heterogeneity in genetic instruments effects, indicating potential violations of the instrumental variable (IV) assumptions, we used the MR Egger intercept test^13^, the Cochran Q heterogeneity test^14^, and the MR pleiotropy residual sum and outlier (MR-PRESSO) test^15^ MR-Egger regression provides test for average pleiotropy: MR Egger regression intercept generally interpreted as average pleiotropic effect across all instruments. MR-Egger has been extended to correct for both measured and unmeasured pleiotropy in MVMR^16^. The Cochran Q test used to identify outliers has been applied in MR to detect average pleiotropy ^15^: pleiotropy can induce heterogeneity of individual ratio estimates^15^. MR-PRESSO detects pleiotropic bias in MR caused by violation of the exclusion restriction IV assumption; extending principal of the Q test, provides a global test to detect pleiotropic bias and identify source of the bias (outlier SNP(s))^15^. MR-PRESSO has been extended to detect pleiotropic bias also in MVMR^15^. MR-PRESSO global tests used to identify outlier SNPS; removing outlier SNPs, we reran MR, and retested so as to determine whether removing outlier(s) resolved detected heterogeneity.

**References.**

1	Karlsson Linnér, R. *et al.* Genome-wide association analyses of risk tolerance and risky behaviors in over 1 million individuals identify hundreds of loci and shared genetic influences. *Nature Genetics* **51**, 245-257, doi:10.1038/s41588-018-0309-3 (2019).

2	Willer, C. J. *et al.* Discovery and refinement of loci associated with lipid levels. *Nat Genet* **45**, 1274-1283, doi:10.1038/ng.2797 (2013).

3	Nikpay, M. *et al.* A comprehensive 1,000 Genomes-based genome-wide association meta-analysis of coronary artery disease. *Nat Genet* **47**, 1121-1130, doi:10.1038/ng.3396 (2015).

4	Malik, R. *et al.* Low-frequency and common genetic variation in ischemic stroke: The METASTROKE collaboration. *Neurology* **86**, 1217-1226, doi:10.1212/wnl.0000000000002528 (2016).

5	Kettunen, J. *et al.* Genome-wide study for circulating metabolites identifies 62 loci and reveals novel systemic effects of LPA. *Nature Communications* **7**, 11122, doi:10.1038/ncomms11122

<https://www.nature.com/articles/ncomms11122#supplementary-information> (2016).

6	Bowden, J., Smith, G. D. & Burgess, S. Mendelian randomization with invalid instruments: effect estimation and bias detection through Egger regression. *Int J Epidemiol* **44**, 512-525, doi:10.1093/ije/dyv080 (2015).

7	Burgess, S., Butterworth, A. & Thompson, S. G. Mendelian Randomization Analysis With Multiple Genetic Variants Using Summarized Data. *Genet Epidemiol* **37**, 658-665, doi:10.1002/gepi.21758 (2013).

8	Burgess, S. & Thompson, S. G. Multivariable Mendelian randomization: the use of pleiotropic genetic variants to estimate causal effects. *American journal of epidemiology* **181**, 251-260, doi:10.1093/aje/kwu283 (2015).

9	Yavorska, O. O. & Burgess, S. MendelianRandomization: an R package for performing Mendelian randomization analyses using summarized data. *Int J Epidemiol* **46**, 1734-1739, doi:10.1093/ije/dyx034 (2017).

10	Davey Smith, G. *et al.* Assessing the suitability of summary data for two-sample Mendelian randomization analyses using MR-Egger regression: the role of the I2 statistic. *Int J Epidemiol* **45**, 1961-1974, doi:10.1093/ije/dyw220 (2016).

11	Bowden, J., Smith, G. D., Haycock, P. C. & Burgess, S. Consistent Estimation in Mendelian Randomization with Some Invalid Instruments Using a Weighted Median Estimator. *Genet Epidemiol* **40**, 304-314, doi:10.1002/gepi.21965 (2016).

12	Hartwig, F. P., Smith, G. D. & Bowden, J. Robust inference in summary data Mendelian randomization via the zero modal pleiotropy assumption. *Int J Epidemiol* **46**, 1985-1998, doi:10.1093/ije/dyx102 (2017).

13	Bowden, J. *et al.* A framework for the investigation of pleiotropy in two-sample summary data Mendelian randomization. *Stat Med* **36**, 1783-1802, doi:10.1002/sim.7221 (2017).

14	Bowden, J. *et al.* Improving the accuracy of two-sample summary-data Mendelian randomization: moving beyond the NOME assumption. doi:10.1093/ije/dyy258 (2018).

15	Verbanck, M., Chen, C. Y., Neale, B. & Do, R. Detection of widespread horizontal pleiotropy in causal relationships inferred from Mendelian randomization between complex traits and diseases (vol 50, 693, 2018). *Nature Genetics* **50**, 1196-1196, doi:10.1038/s41588-018-0164-2 (2018).

16	Rees, J. M. B., Wood, A. M. & Burgess, S. Extending the MR-Egger method for multivariable Mendelian randomization to correct for both measured and unmeasured pleiotropy. *Stat Med* **36**, 4705-4718, doi:10.1002/sim.7492 (2017).
